# omicsMIC: a Comprehensive Benchmarking Platform for Robust Comparison of Imputation Methods in Mass Spectrometry-based Omics Data

**DOI:** 10.1101/2023.09.12.557189

**Authors:** Weiqiang Lin, Jiadong Ji, Kuan-Jui Su, Chuan Qiu, Qing Tian, Lan-Juan Zhao, Zhe Luo, Hui Shen, Chong Wu, Hongwen Deng

**Affiliations:** Tulane Center for Biomedical Informatics and Genomics, Deming Department of Medicine, School of Medicine, Tulane University, New Orleans, LA 70112, USA; Institute for Financial Studies, Shandong University, Jinan 250100, China; Department of Biostatistics, The University of Texas MD Anderson Cancer Center, Houston, TX, USA

## Abstract

Mass spectrometry is a powerful and widely used tool for generating proteomics, lipidomics, and metabolomics profiles, which is pivotal for elucidating biological processes and identifying biomarkers. However, missing values in spectrometry-based omics data may pose a critical challenge for the comprehensive identification of biomarkers and elucidation of the biological processes underlying human complex disorders. To alleviate this issue, various imputation methods for mass spectrometry-based omics data have been developed. However, a comprehensive and systematic comparison of these imputation methods is still lacking, and researchers are frequently confronted with a multitude of options without a clear rationale for method selection. To address this pressing need, we developed omicsMIC (mass spectrometrybased *omics* with Missing values Imputation methods Comparison platform), an interactive platform that provides researchers with a versatile framework to simulate and evaluate the performance of 28 diverse imputation methods. omicsMIC offers a nuanced perspective, acknowledging the inherent heterogeneity in biological data and the unique attributes of each dataset. Our platform empowers researchers to make data-driven decisions in imputation method selection based on real-time visualizations of the outcomes associated with different imputation strategies. The comprehensive benchmarking and versatility of omicsMIC make it a valuable tool for the scientific community engaged in mass spectrometry-based omics research. OmicsMIC is freely available at https://github.com/WQLin8/omicsMIC.

## Introduction

The recent advancements in mass spectrometry-based omics, such as proteomics, lipidomics, and metabolomics, have ushered in a new era of scientific discovery, advancing our understanding of biological mechanisms (1) and potentially reshaping the discovery of biomarkers (2-5), drug discovery and precision medicine (6,7) for human complex disorders by providing insights into molecular biology and enabling comprehensive molecular analysis. Especially, the integration of other omics data with mass spectrometry-based omics data has emerged as a potent strategy for advancing our comprehension of complex biological progress (8,9).

However, one of the main drawbacks of mass spectrometry-based omics data is that it typically contains a large proportion of missing values, even in the range of 30-50% (10,11). Commonly, these missing values can be attributed to either the genuine absence of the compound in the measured sample or the presence of the molecular feature at a concentration below the mass spectrometer’s detection limit. Therefore, many studies handled missing values with simple value replacement methods like zero (12-14), half of the min value (15-18), and the min value (19-21) of the corresponding feature. However, the issue of missing data is a complex problem that can arise in various situations including: (1) Mass spectrometry techniques may encounter technical problems during data acquisition, such as instrument errors, signal interference, or instrument malfunctions (22-26); (2) Prior to mass spectrometry analysis, samples undergo a series of preparation and extraction steps. These steps may involve chemical reactions, sample handling, or extraction processes, which introduce variability and uncertainty (22-25,27); (3) Processing and interpreting mass spectrometry data is a complex process that involves steps such as signal denoising, background correction, and quality filtering. During the processing, certain data points may not meet specific quality standards or the limitations of the algorithms. These issues can result in certain data points not being accurately recorded or obtained and thus being treated as missing data (28,29). Inadequately addressing these missing data can introduce bias in the subsequent statistical analysis and interpretation of mass spectrometry-based data, potentially compromising the reliability of the downstream analysis and the accuracy of the results.

Imputation is a common approach to dealing with missing data, which treats missing values using the information that is available from the existing data. So far, numerous imputation methods (30-37) have been proposed for handling missing values in -omics studies. In this study, we roughly divide incorporated imputation methods into three categories: (i) simple value replacement, (ii) model-based approaches, and (iii) machine learning-based approaches. Simple value replacement is a commonly used technique for handling missing data in various fields like zero, half of the minimum value, and minimum value. This strategy can quickly fill in missing values and allow for downstream analyses to be conducted on complete datasets. However, it may skew the distribution or underestimate measures of variance and lead to more bias (29,38). To account for this limitation of simple value replacement, many model-based imputation methods have been developed, which leverage statistical and computational models to estimate missing values, taking into account the patterns and relationships present in the data. Examples include Bayesian PCA (BPCA) (34) and Singular Value Decomposition (SVD) imputation (35). Furthermore, machine learning-based approaches have become increasingly popular, as they can handle diverse data distributions and complex relationships. For instance, K-nearest neighbors (KNN) imputations (36) use similarity between samples to predict missing values and random forest-based methods (30,37) use the random forest ensemble algorithm to make predictions.

Numerous imputation methods have been proposed and previous studies suggested that different methods may be required to achieve good performance under different circumstances (28,29,39,40). However, a comprehensive and systemic comparison of the performance of these imputation methods under different conditions is still lacking. In this study, we develop omicsMIC (mass spectrometry-based *omics* with Missing values Imputation methods Comparison platform), a user-friendly platform that provides a versatile framework for simulating and evaluating a diverse range of imputation strategies, tailored to users’ specific datasets. Our platform can help users determine the most appropriate imputation method for their datasets with specific objectives and allow users to perform the imputation on their datasets once the preferred imputation approach is determined. We anticipate that this platform will be helpful for the community, particularly for researchers without an extensive background in computer science or programming within this biomedical field.

### Overview of omicsMIC interactive server

omicsMIC is a comprehensive application that allows advanced comparison of various imputation methods for mass spectrometry-based omics data in a user-interactive fashion. The main interface of omicsMIC displays the analysis steps on an expandable menu on the left side and shows a guidance section on the right side (**Figure 1**). The analysis steps include Upload data file, Data quality and control, Data simulation, and Data imputation. The whole workflow of the omicsMIC is shown in **Figure 2** and is described below with available options.

**Figure 1.**
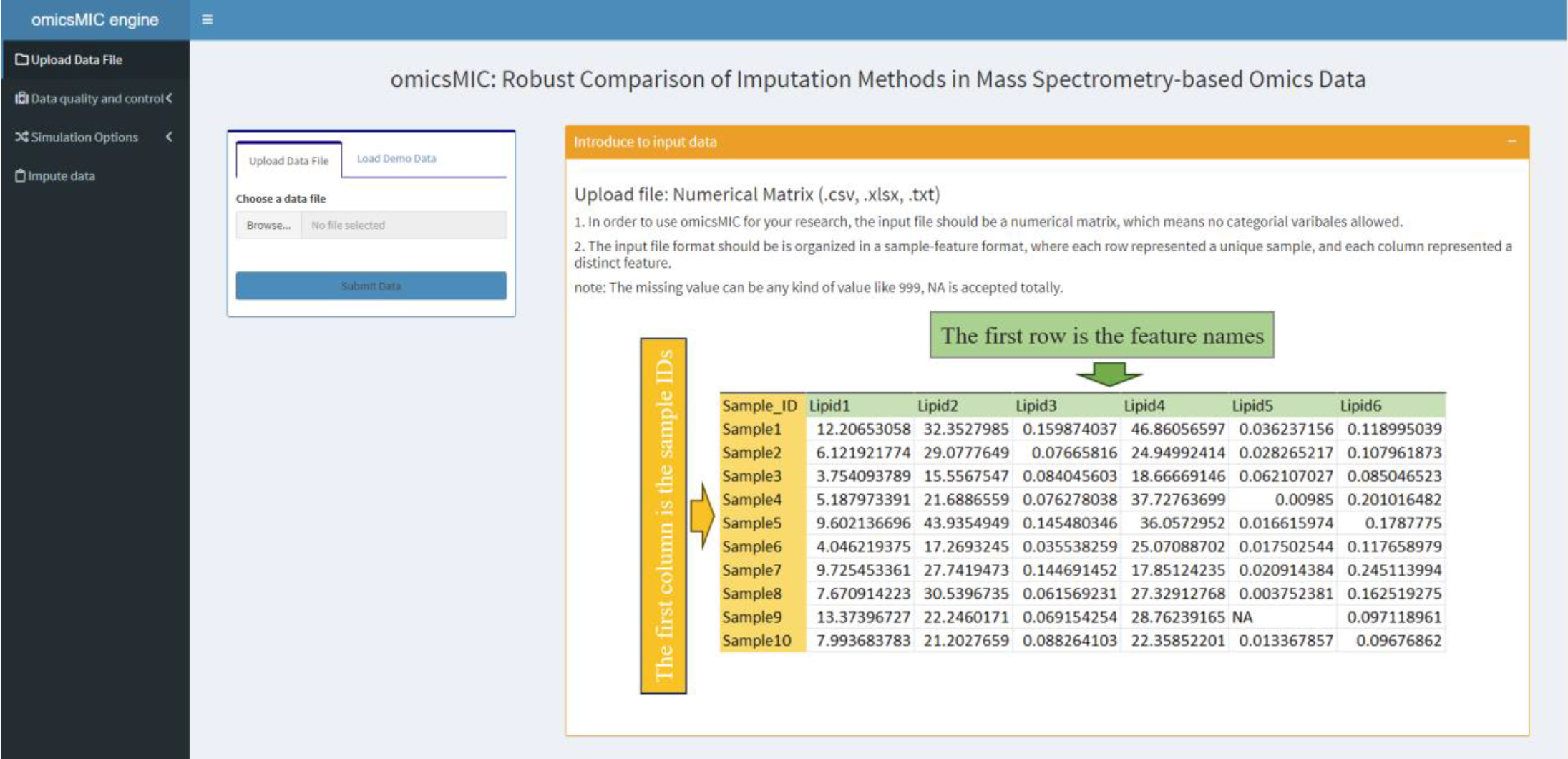
OmicsMIC main interface.

**Figure 2.**
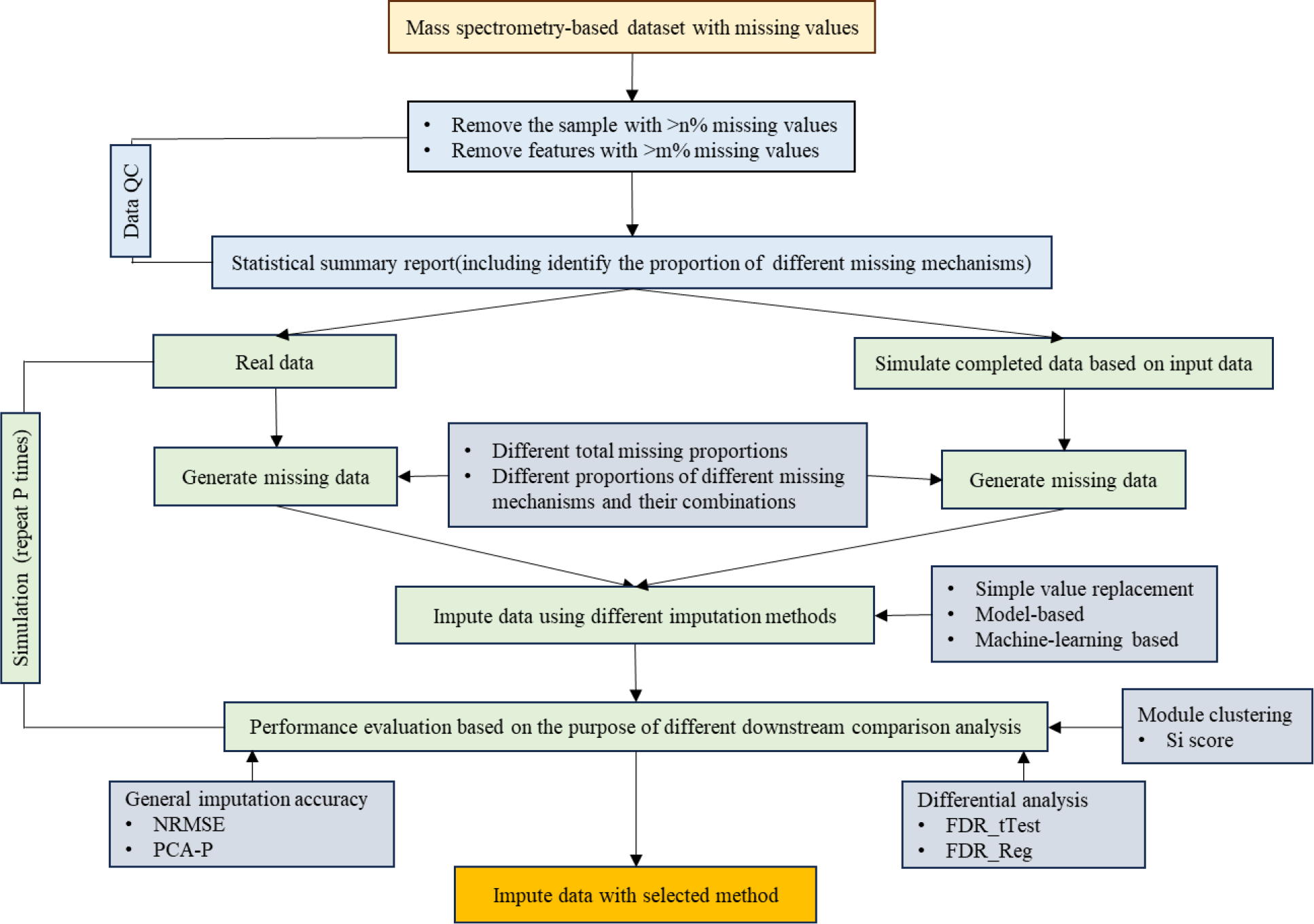
The workflow of OmicsMIC.

### Input

OmicsMIC analysis starts with uploading a data file (**Figure 3A**), containing mass spectrometrybased quantitative data. The file should be organized in a sample-feature format, where each row represents a unique sample, and each column represents a distinct feature. OmicsMIC can accommodate various file formats, including CSV, and XLSX. Once the data is uploaded, the program will provide a quick overview of the uploaded dataset, allowing users to verify that the correct data has been uploaded (**Figure 3B**). We also provide a demo dataset from an empirical lipidomics study to help users familiarize themselves with the platform’s functionalities the platform (**Figure 3C**).

**Figure 3.**
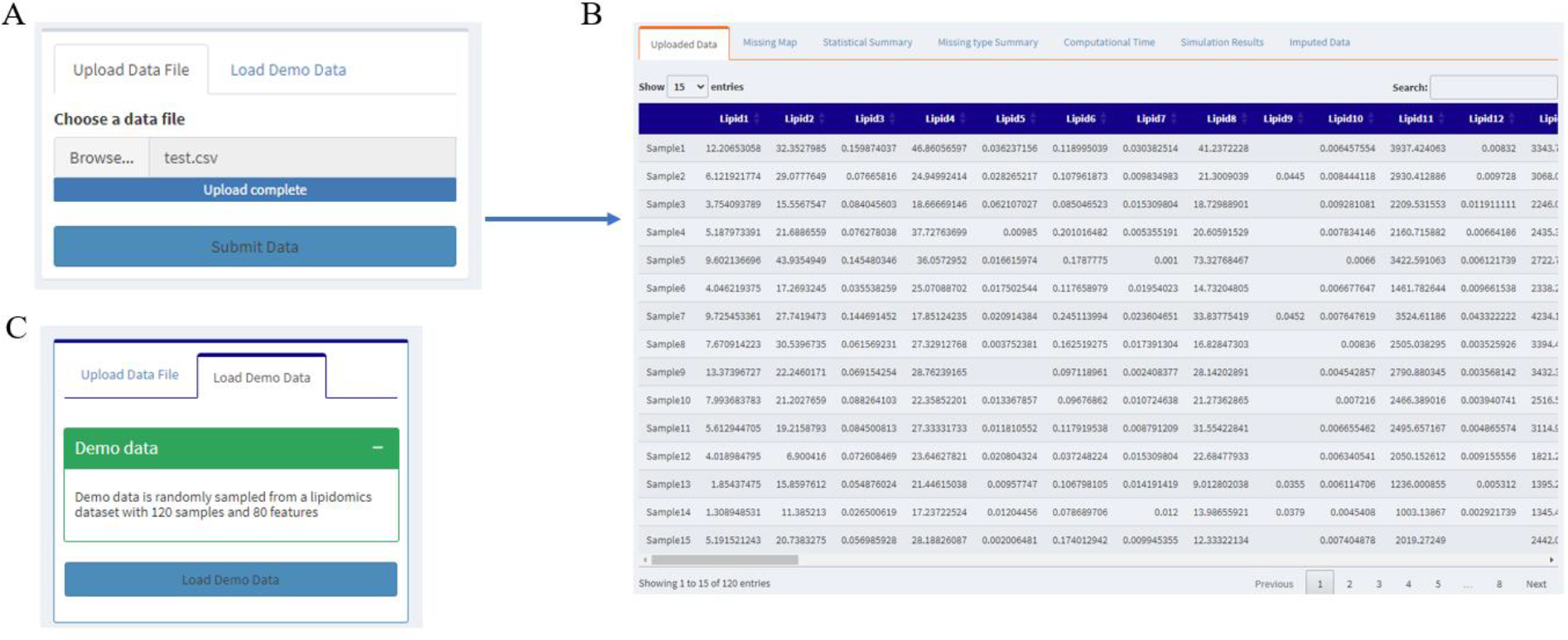
Data input interfaces. (A) A quantitative dataset which is organized in a sample-feature format is required. (B) The uploaded data is shown correspondingly. (C) Demo dataset for users to get familiar with omicsMIC.

### Data quality and control

Within the omicsMIC platform, two vital data quality control functionalities are implemented, allowing users to tailor their data preparation according to their specific requirements.

1. *Removal of High Missing Data:* Our platform allows users to control data quality precisely by customizing the threshold for removing samples and features with excessive missing data. Users can specify missing rate thresholds (**Figure 4A**) so that samples and features with missing rates over the thresholds will be excluded from the subsequent analyses. Upon userdefined criteria, omicsMIC will generate a missing data pattern heatmap (**Figure 4B**), visually representing the distribution of missing values across samples and features. This heatmap offers valuable insights into the data’s completeness and assists users in identifying patterns or trends in missing data.
2. *Statistical Summary:* In this section, we present a comprehensive statistical summary after the removal of samples and features surpassing the specified missing data threshold, providing users with essential insights into the characteristics of their dataset alongside the statistical metrics and missing mechanisms. Users have the flexibility to select correlation methods and goodness of fit indices (**Figure 4C**), tailoring the analysis to their specific needs. For each variable in the dataset, omicsMIC will calculate and report the following statistical metrics: missing number, missing %, min, mean, max, variance, 25th percentile, 50th percentile, 75th percentile, missingness type, and distribution (**Figure 4D**). Additionally, it will also provide the user with a summary report of the proportion of different missing mechanisms (**Figure 4E**).

**Figure 4.**
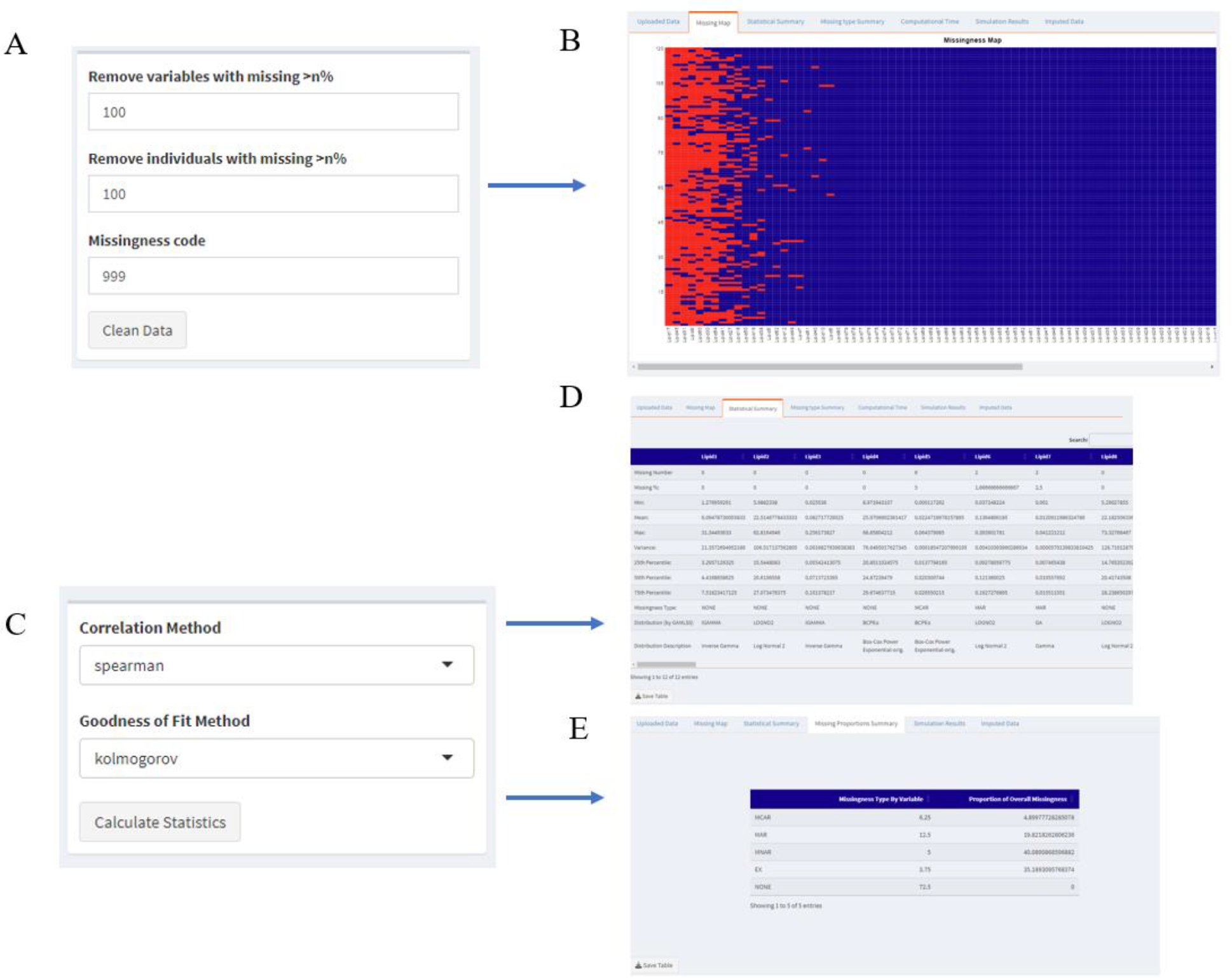
Data quality and control interfaces. omicsMIC allows users to select different processing and analysis options and assess the characteristics of their data with visualized output. (A) The parameters for setting thresholds of removing samples and features. (B) Missing value heatmap after removing high missing proportion features or samples. (C) The parameters for data statistical summary report. (D) The data summary report. (E) The summary report of different missing mechanisms.

### Data simulation

The omicsMIC platform introduces a potent data simulation feature, armed with seven customizable parameters. These parameters have been thoughtfully crafted to equip users with a versatile toolkit for effectively appraising various imputation methods. Here, we describe these parameters in detail:

1. *Simulate Data or Not:* This parameter empowers users to decide whether they wish to generate simulated data based on the characteristics of their uploaded dataset for imputation method assessment. If users want to use real data to do downstream simulations, the missing value in the uploaded dataset will be excluded by removing samples and features to get a completed dataset (**Figure 5A**).
2. *Simulation Iterations:* Users can define the number of simulation iterations, a pivotal factor for robust evaluations. Commonly, multiple iterations introduce diversity into the simulations and thus increase the reliability of the results (**Figure 5B**).
3. *Missingness Simulation Iterations:* Users can adjust the parameter for missing data generation, which specifies the number of missing data instances created in each simulation (**Figure 5C**).
4. *Missingness Percentage:* Users can predefine the overall missingness proportion in the simulated dataset (**Figure 5D**). The ability to vary this parameter enables exploration of different levels of data incompleteness during simulations, a fundamental aspect of method assessment. It can provide users with the effects of different imputation methods under various missing proportions, helping users decide on the appropriate missing proportion for imputation and downstream data analysis. The whole data missing proportion results from the Data quality and control step will be incorporated as a scenario into the simulation automatically.
5. *Missing type ratio:* This parameter defines different missing mechanism ratio settings (**Figure 5E**), pivotal for evaluating imputation method stability under diverse missing mechanisms. In this study, we consider three types of missing mechanisms: missing not at random (MNAR), missing at random (MAR), and missing completely at random (MCAR) (41). The different missing mechanism ratios calculated from the data quality and control step will be included as a scenario for the simulation automatically.
6. *Imputation Methods:* In the omicsMIC application, a total of 28 imputation methods are included. Imputation methods are categorized into simple value replacement, model-based, and machine learning-based techniques, offering users flexibility tailored to their research needs (**Table 1, Figure 5F-H**). omicsMIC also provides the estimated computational time for each method to assist users in aligning with time constraints (**Figure 6**), which was obtained using the platform’s default parameters with demo data.
7. *Evaluation Methods:* omicsMIC offers various evaluation metrics for assessing imputation methods. Normalize root means standard error (NRMSE) (32) and PCA-Procrustes analysis (PCA-P) (29) are applied to evaluate the overall difference and the alternation in the distribution pattern between the original and the imputed dataset. To evaluate the potential effects of imputation on differential analysis, the False Discovery Rate is performed to compare the ability of different imputation methods to correctly recover variables with differences in the imputed data compared to the original data. Considering the practical requirements for real-world applications, we evaluate the performance of different imputation methods on these two metrics based on t-tests (FDR_tTest) and regression analyses (FDR_Reg). To evaluate the effects of imputation on clustering analysis, the mean Silhouette coefficient (Si score) (42) is used to evaluate the clustering performance of various imputation methods that combine cohesion and separation and can compare the quality of clustering outcomes obtained from different imputation techniques. Different evaluation metrics will help guide users to select methods that best suit their research objectives (**Figure 5I**).

**Figure 5.**
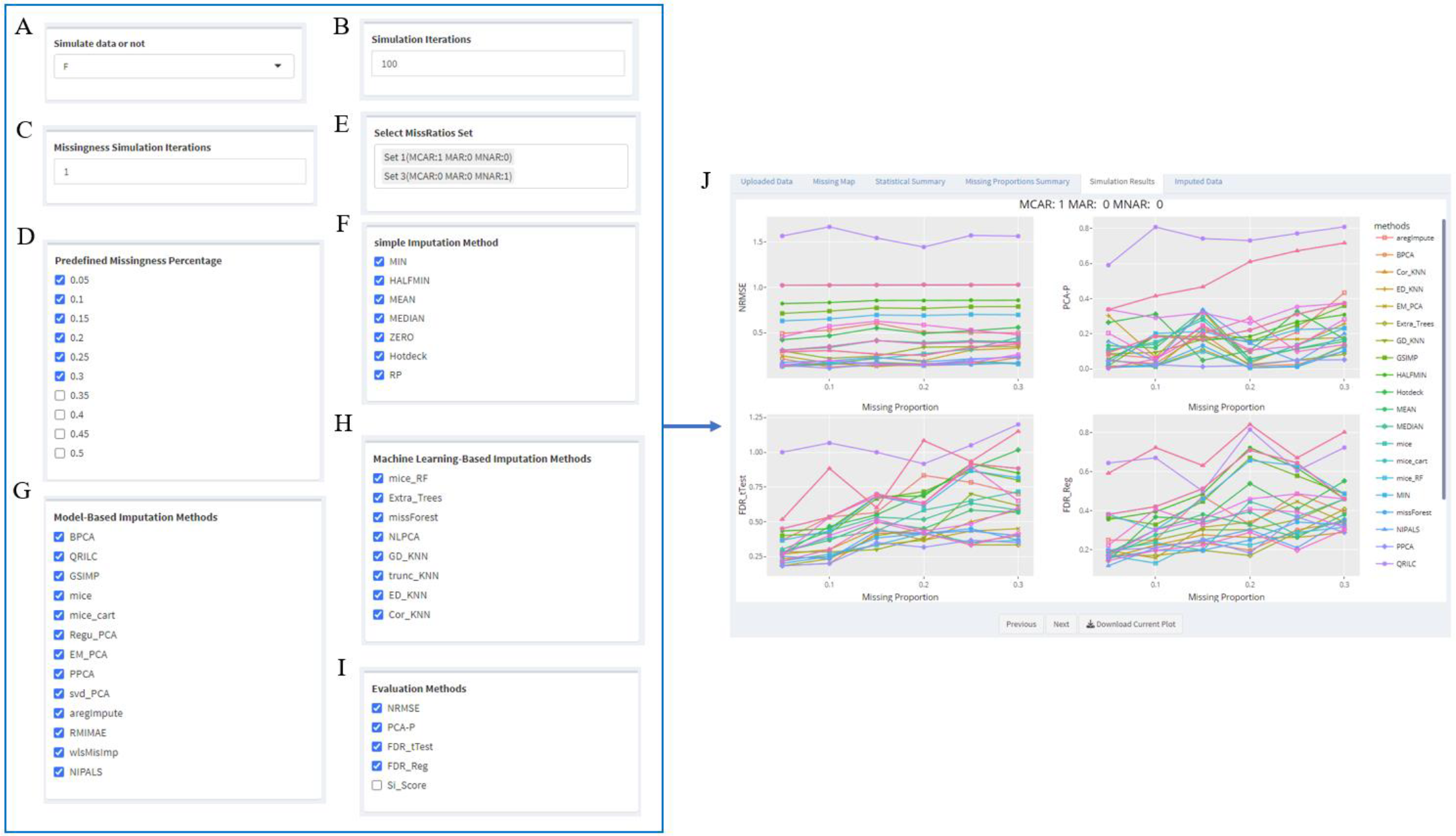
Simulation interfaces. omicsMIC empowers users to actively engage in the assessment and comparison of different imputation methods for their data. Through intuitive user controls, individuals can set simulation parameters and execute imputation methods, all while benefiting from real-time, interactive visualization to evaluate the results. Once the preferred imputation approach is determined, users can seamlessly export static plots for inclusion in publications and reports.

**Table 1.** Brief description of imputation methods evaluated in omicsMIC.

| Category | Methods | Source | Reference |
| --- | --- | --- | --- |
| Simple value replacement | Zero |  |  |
|  | Half-min |  |  |
|  | Min |  |  |
|  | Mean |  |  |
|  | Median |  |  |
|  | hotdeck | VIM | (43) |
|  | Random replacement |  |  |
| Model-based imputation method | BPCA | pcaMethods | https://github.com/hredestig/pcamethods |
|  | QRILC | imputeLCMD | https://github.com/cran/imputeLCMD |
|  | GSimp | https://github.com/WandeRum/GSimp | (33) |
|  | mice | mice | (44) |
|  | mice_cart | mice | (44) |
|  | svd_PCA | pcaMethods | https://github.com/hredestig/pcamethods |
|  | Regular_PCA | missMDA | (45) |
|  | EM_PCA | missMDA | (45) |
|  | PPCA | pcaMethods | https://github.com/hredestig/pcamethods |
|  | aregImpute | Hmisc | https://github.com/harrelfe/Hmisc |
|  | rmiMAE | https://github.com/NishithPaul/missingImputation/blob/main/rmiMAE.R | (31) |
|  | wlsMisImp | https://github.com/NishithPaul/tWLSA | https://github.com/NishithPaul/tWLSA |
|  | NIPALS | pcaMethods | https://github.com/hredestig/pcamethods |
| Machine learning-based imputation methods | mice_RF | missRanger | https://github.com/mayer79/missRanger |
|  | Extra_Trees | missRanger | https://github.com/mayer79/missRanger |
|  | missForest | missForest | (30) |
|  | NLPCA | pcaMethods | https://github.com/hredestig/pcamethods |
|  | GD_KNN | VIM | (43) |
|  | Cor_KNN | MmPalateMiRNA | (46) |
|  | trunc_KNN | https://github.com/WandeRum/GSimp | (33) |
|  | ED_KNN | https://github.com/WandeRum/GSimp | (33) |

**Figure 6.**
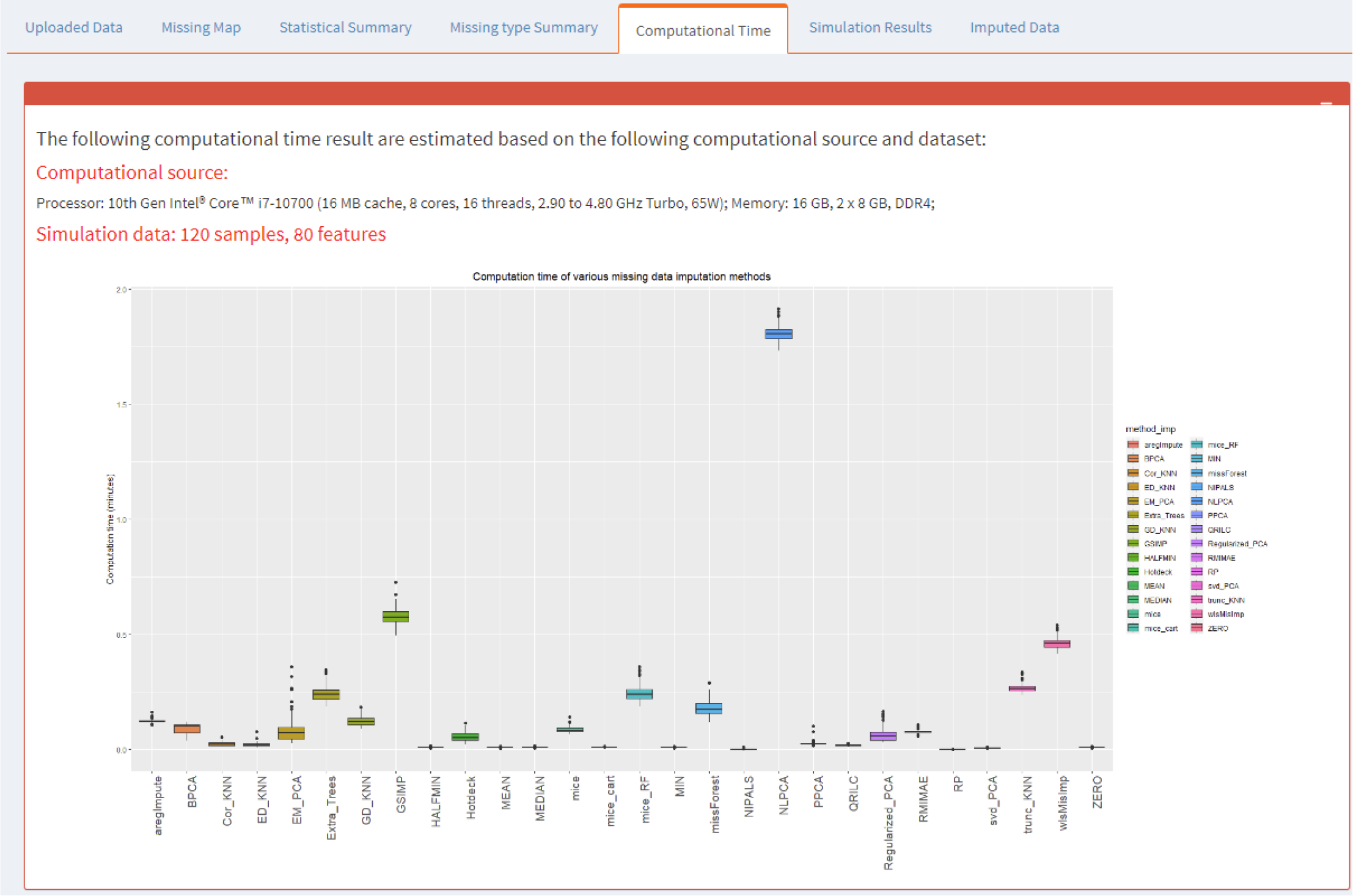
Boxplot of Average Execution Times for Various Missing Data Imputation Methods.

The “Data Simulation” section presents unparalleled flexibility and control for imputation method assessment. Leveraging these parameters, researchers can conduct rigorous evaluations, make informed decisions, and select the most appropriate imputation methods, thereby enhancing the quality and trustworthiness of their data analysis. The comparison results are presented as interactive figures which can be exported as PNG files (**Figure 5J**).

### Data imputation

After selecting an appropriate method for imputing missing data, the users can use omicsMIC to perform the imputation on their uploaded datasets and download the imputed dataset for further downstream data analysis (**Figure 7**).

**Figure 7.**
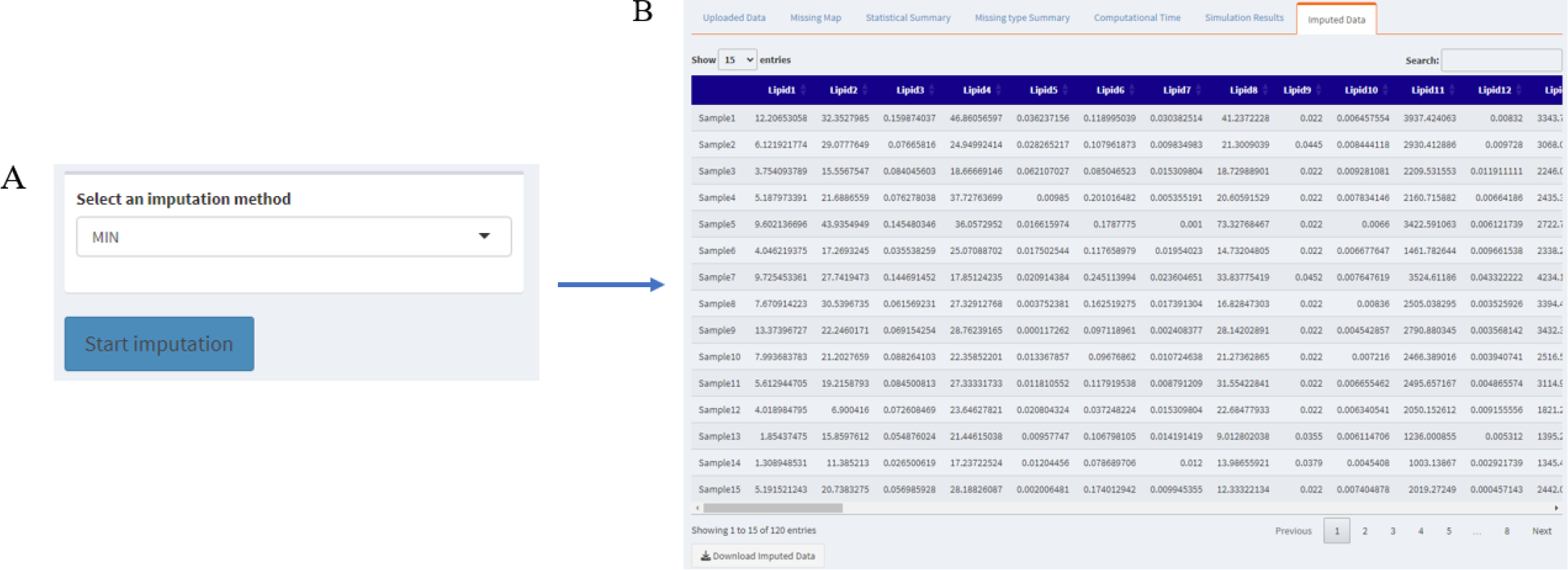
Imputation interfaces. The imputation function is provided by omicsMIC and the user can download the imputed data.

## Conclusion

omicsMIC offers a versatile framework for simulating and evaluating a wide array of imputation strategies for mass spectrometry-based omics data. Given the inherent heterogeneity of biological data, omicsMIC equips users with real-time visualizations of imputation outcomes, facilitating informed and rational method selection. Notably, most imputation strategies incorporated in the omicsMIC platform are not only for mass spectrometry-based omics data but can also be applied to other types of continuous data. Furthermore, as the source code of omicsMIC will be publicly shared, the platform has the potential for future updates to accommodate newly developed methods and additional evaluation criteria. Therefore, omicsMIC is a valuable tool for the development and evaluation of novel imputation methods.

## Funding

This work is partially supported by grants from the NIH (U19AG055373, R01AG061917, R01AR069055).

## Conflict of interest statement

None declared.

## Notes

### Competing Interest Statement

The authors have declared no competing interest.

